# RNA-Chrom: a manually-curated analytical database of RNA–chromatin interactome

**DOI:** 10.1101/2022.12.10.519346

**Authors:** G. K. Ryabykh, S. V. Kuznetsov, Y. D Korostelev, A. I. Sigorskikh, A. A. Zharikova, A. A. Mironov

**Affiliations:** Faculty of Bioengineering and Bioinformatics, Lomonosov Moscow State University, Leninskiye Gory, 119234, Moscow, Russia; Kharkevich Institute for Information Transmission Problems RAS, Bolshoy Karetny per., 127051, Moscow, Russia; National Medical Research Center for Therapy and Preventive Medicine, Petroverigsky per., 101000, Moscow, Russia

**Keywords:** non-coding RNAs, RNA–chromatin interactome, contact maps, database

## Abstract

Every year there is more and more evidence that non-coding RNAs play an important role in biological processes affecting various levels of organisation of living systems: from the cellular (regulation of gene expression, remodeling and maintenance of chromatin structure, co-transcriptional suppression of transposons, splicing, post-transcriptional RNA modifications, etc.), to cell populations and even organismal ones (development, aging, cancer, cardiovascular and many other diseases). The development and creation of mutually complementary databases that will aggregate, unify and structure different types of data can help to reach the system-level of studying non-coding RNAs. Here we present the RNA-Chrom manually-curated analytical database, which contains the coordinates of billions of contacts of thousands of human and mouse RNAs with chromatin. Through the user-friendly web interface (https://rnachrom2.bioinf.fbb.msu.ru/), two approaches to the analysis of the RNA–chromatin interactome were implemented. Firstly, to find out whether the RNA of interest to a user contacts with chromatin, and if so, with which genes or DNA loci? Secondly, to find out which RNAs are in contact with the DNA locus of interest to a user (and probably participate in its regulation), and if there are such, what is the nature of their interaction? For a more detailed study of contact maps and their comparison with other data, the web interface allows a user to view them in the UCSC Genome Browser.

## Introduction

Back in the 1960s, it was found that a large amount of different RNAs is associated with chromatin [1, 2]. However, it remains unknown what kind of RNAs they are, which chromatin loci they prefer to interact with, and what function they perform there. Much later, mainly using molecular biochemical methods, the functions of some non-coding RNAs were determined: XIST, which is responsible for X-chromosome dosage compensation in mammals [3], Kcnq1ot1, which is involved in imprinting [4], and others. With the development of new methods, especially those involving a high-throughput sequencing step, more and more is becoming known about new chromatin-associated RNAs and their mechanisms of action.

For example, roX1 and roX2 RNAs are responsible for X-chromosome dosage compensation in *Drosophila*. There is also a number of regulatory RNAs (XIST, HOTAIR, MEG3, Paupar, ANRIL, TERRA, SRA, etc.) affect gene expression by attracting such chromatin modifiers as TrxG and PRC1/PRC2 to certain loci. Another example is MALAT1 and NEAT1, which are associated with such nuclear structures as speckles and paraspeckles and regulate gene expression. Moreover, Firre can serve as a local organising factor to ensure a topological proximity of trans-sites and its genomic locus; lnc-NR2F1 is involved in neurogenesis; Bloodlinc is involved in erythropoiesis; DACOR1 interacts with maintaining DNA methyltransferase 1 and is expressed at a higher level in normal cells of the healthy colon and at a lower level in colon cancer cell clones. Many more biological examples of the interaction of RNA with chromatin can be cited [5].

To identify DNA loci contacted by non-coding RNAs, a number of experimental methods have been developed that can be divided into two groups. The first group of methods (RAP [6], CHART-seq [7], ChIRP-seq [8], dChIRP-seq [9], ChOP-seq [10], and CHIRT-seq [11]) solves the problem of finding contacts of a predetermined RNA. We will call this group of methods “one RNA to all DNA loci”, or “one-to-all”. The main idea of all these methods is as follows. Cells are fixed, resulting in covalent crosslinking of macromolecules. Next, DNA is fragmented, which results in a mixture of various complexes, including the RNA of interest with genomic DNA. Biotinylated oligonucleotides complementary to RNA of interest are used to isolate specific complexes. After isolation of the complexes with the target RNA on the streptavidin beads, the complexes are cleaved, the protein fraction is removed, and the DNA is sequenced. It is assumed that the target RNA interacts with these DNA loci. The disadvantage of this approach is that the target RNA must be determined in advance [5].

Another group of methods is designed to search for all possible DNA and RNA contacts in a cell (MARGI [12], GRID-seq [13, 14, 15], ChAR-seq [16], iMARGI [17, 18], RADICL-seq [19], Red-C [20]). We will call this group of methods “all RNAs to all DNA loci”, or “all-to-all”. The main idea of this approach is that after cell fixation and DNA fragmentation, proximity ligation is carried out using a specially designed bivalent biotenilated bridge. After ligation of the bridge to RNA and then to DNA and reverse transcription, the chimeric constructs are sequenced. The result is a set of contacts of various RNAs and DNA loci (Figure 1). The disadvantage of this approach is that a large number of reads is needed to obtain reasonable data [5].

**Fig. 1.**
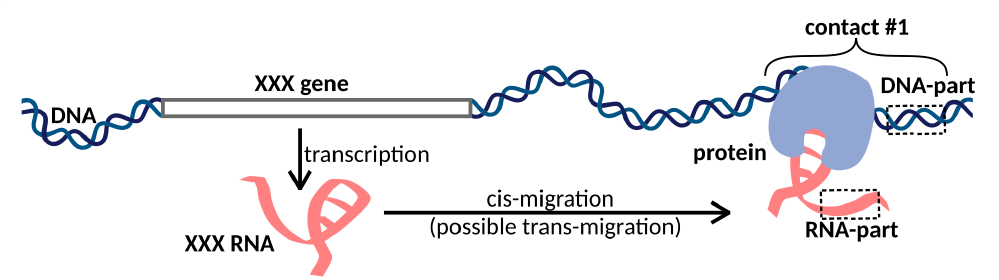
XXX RNA interacts with the DNA locus and forms contact #1. In the case of one-to-all methods, we see only the DNA-parts of the contacts, while in the case of all-to-all methods, we see both DNA-parts and RNA-parts of the contacts. Cis- and trans-migration is the migration of RNA within and outside the parent chromosome, respectively.

There are many databases available that facilitate the system-level in the study of the non-coding RNAs action mechanisms [21]. They can roughly be divided into “general” databases, which aggregate a variety of non-coding RNA data (e.g. NONCODEV5 [22]), and “highly specialised” ones, which focus either on a biological process (e.g. cancer – Cancer lncRNA Census [23]), or on a biological system (e.g. the cardiovascular system [24]), or on a specific type of data (e.g. histone modification and transcriptome data – HiMoRNA [25]). In the case of the RNA–chromatin interactions, there is no database that is highly specialised on this type of data. On the other hand, there are a number of general databases with a web interface, which, among other things, contain collections of RNA–DNA contacts: RNAInter [26] and LnChrom [27] (unfortunately, the LnChrom has not been available for more than a year). RNAInter is the most comprehensive resource on the RNA interactome. An elementary object of this database is an RNA contact confirmed in one or another experiment, for which confidence score is calculated. However, this resource contains a small number of genome-wide experiments on RNA– chromatin contacts, in particular, apart from MARGI, there are no all-to-all data.

Comparative analysis of the RNA–chromatin interactome is of great scientific interest. To solve this problem, we have developed a highly specialised analytical database (https://rnachrom2.bioinf.fbb.msu.ru/) that contains all available genome-wide RNA–DNA interactions data. Since there is no standard protocol for processing these data, it is difficult to conduct a comparative analysis of RNA–chromatin contacts. Here we have standardised the data processing protocol and implemented it starting with raw reads. The RNA-Chrom database allows a user not only to download data processed by a single protocol, but also to perform various methods of data analysis and comparison in real time. It is also possible to view contact maps in the UCSC Genome Browser [28] to study them in more detail and compare them with other data, such as DNA methylation data.

## Materials and methods

### General scheme of a web application

“Front-end” was developed using “Node.js”^1^ (an asynchronous event-driven JavaScript runtime), “React.js”^2^ and “Redux”^3^ libraries and implemented as a “Single Page Application”. The “Material-UI V4”^4^ library was taken as the basis for the web interface elements, and the “Plotly JavaScript Open Source Graphing Library”^5^ was taken to create interactive plots. “Back-end” was implemented using the Python web microframework “Quart”^6^ as it supports asynchronous database requests. The database itself, which stores RNA-chromatin contacts, was created on the basis of “ClickHouse”^7^ (the Open Source OLAP database management system), due to which a user’s waiting time for any of their analytical requests was reduced to seconds.

### Extraction of the RNA–chromatin interactions data

Since the first articles with all-to-all methods appeared only in 2017, it was not difficult to find data corresponding to them. Things were quite different with one-to-all data. We first searched for them in Gene Expression Omnibus^8^ using the keywords “RAP-seq”, “CHART-seq”, “ChIRP-seq”, “dChIRP-seq”, “ChOP-seq”, “CHIRT-seq” and taking into account only human and mouse data sets. Then we went as far as possible through articles that referred to the main one-to-all methods (RAP [6], CHART-seq [7], ChIRP-seq [8], dChIRP-seq [9], ChOP-seq [10] and CHIRT-seq [11]). Surprisingly, we found a large number of publications that used one-to-all methods, but there was no publicly available data. Only one author responded to our request and provided us with RAP-data for Firre RNA [29]. A total of 66 articles were found that had data in the public domain.

### Universal data processing protocol

There are many approaches to RNA–chromatin interactions data processing, but each of them is tailored to the data obtained by a certain experimental method. Since it is necessary to use a single protocol for data unification and further comparative analysis, the authors of the LnChrom database used the protocol from the ChIRP article [8]. Our database contains both all-to-all and one-to-all data, and we will base on the protocol applied in the Red-C experiment [20] (Figure 2), the details of which are disclosed in the Supplementary Text 1.

**Fig. 2.**
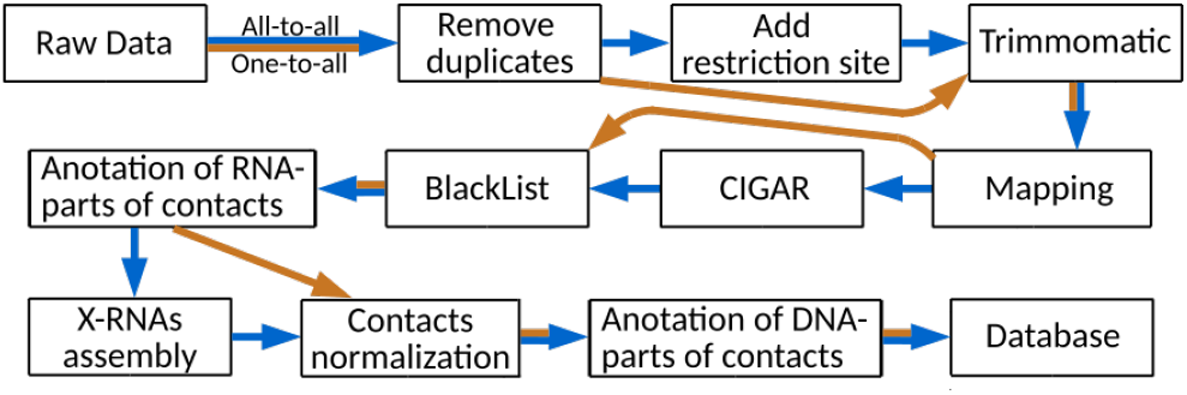
RNA–chromatin interactions data processing protocol. Blue arrows correspond to all-to-all data processing steps, orange arrows are related to one-to-all data.

Raw data was downloaded from Gene Expression Omnibus and European Nucleotide Archive^9^. Possible PCR duplicates were removed via FastUniq [30] and SeqKit [31] tools. “Add restriction site” step was performed only for all-to-all data in strict accordance with the recommendations of the original articles. For all data, we used TRIMMOMATIC (v0.39) [32] for detection of low-quality position in each forward and reverse read. One-to-all and all-to-all data were mapped to the canonical chromosomes of the human and mouse genomes (GRCh38 and GRCm38 assemblies, respectively) with HISAT2 program (version 2.1.0) [33]. After that, the orientations of the RNA-parts of the contacts were refined (Supplementary Text 2). It turned out that in the MARGI experiment [12], in most cases a random strand was read, and the orientations of the RNA-parts were lost. Based on this finding, it was decided to exclude these data sets from any further analysis. To search for and process reads with splicing corresponding to RNA-parts of the contacts, the mapping information presented in the CIGAR field was analysed. In turn, in accordance with the RADICL-seq protocol [19], those DNA-parts of the contacts that fell into the regions from the ENCODE BlackList were removed.

It is important for any contact to know the source gene. To do this, we collected the general gene annotation (Tables 1 and 2), balancing between its large size and the low representation of certain types of RNA. The clusters of unannotated RNA-parts of contacts were found and they were named X-RNAs. Only the contacts with RNA-parts that intersect the genes from the general annotation were added to the database. The others were named “Singletons” and were not used. Having passed the data through all the previous steps of the protocol, we got the final number of contacts for each experiment. For all-to-all and one-to-all experiments, a background model was calculated, according to which each contact (in addition to the original or “Raw” single value) was assigned a “Normalized” value. Two additional normalizations were obtained for the one-to-all data: “Norm. & in peaks” (background-normalized contacts crossing MACS2 peaks) and “Raw & in peaks” (not-normalized contacts crossing MACS2 peaks). It is important to note that the median number of “reads in peaks” for all experiments is 30 times less than the median number of reads that have passed all processing steps (Supplementary Figure 1). Thus, in our database there are 4 types of normalization. As a final step before loading the data into the database, we annotated DNA-parts of the contacts with genes and near-gene regions. To get the comparable characteristic of RNA contactability in all-to-all experiments we introduced a “CPKM” metrics – Contacts Per Kilobase of RNA length per Million filtered contacts in the experiment.

**Table 1.**
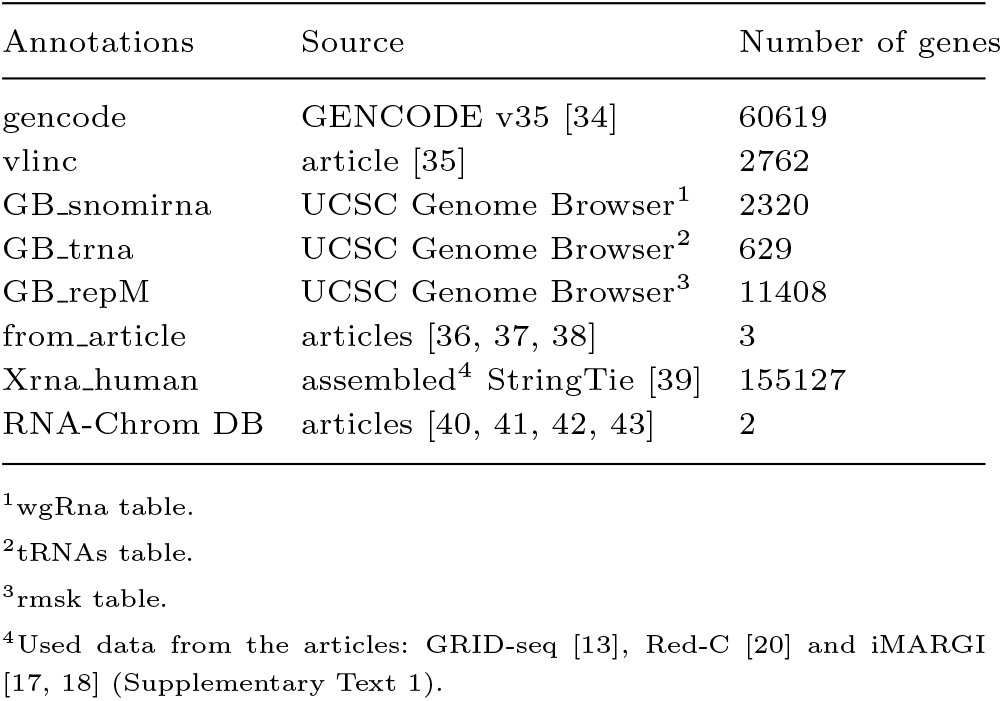
Human gene annotations (only from canonical chromosomes)

**Table 2.** Mouse gene annotations (only from canonical chromosomes)

| Annotations | Source | Number of genes |
| --- | --- | --- |
| gencode | GENCODE M25 [34] | 55364 |
| GB_trna | UCSC Genome Browser<sup>1</sup> | 434 |
| GB_repM | UCSC Genome Browser<sup>2</sup> | 18770 |
| from_article | articles [44, 45, 46, 47] | 4 |
| Xrna_mouse | assembled<sup>3</sup> StringTie [39] | 14333 |
| RNA-Chrom DB | articles<sup>4</sup> | 9 |

<sup>1</sup>tRNAs table.
<sup>2</sup>rmsk table.
<sup>3</sup>Used data from the articles: GRID-seq [13], RADICL-seq [19] (Supplementary Text 1).
<sup>4</sup>[11, 41, 48, 49, 50, 51, 52, 53, 54, 55, 56].

According to the summary statistics (Supplementary Table 1), for all-to-all data compared to one-to-all data, the largest number of reads is filtered out in the “Mapping” step (Figure 3). This is because for all-to-all data, we require that the RNA- and DNA-parts of each contact are both correctly mapped, otherwise they will be filtered out. As for the one-to-all data, there are several data sets among them that are not credible. For example, GSM3073889 and GSM3073888 (human lincDUSP RNA) have less than 4000 raw reads and no MACS2 peaks. However, they are still in the database.

**Fig. 3.**
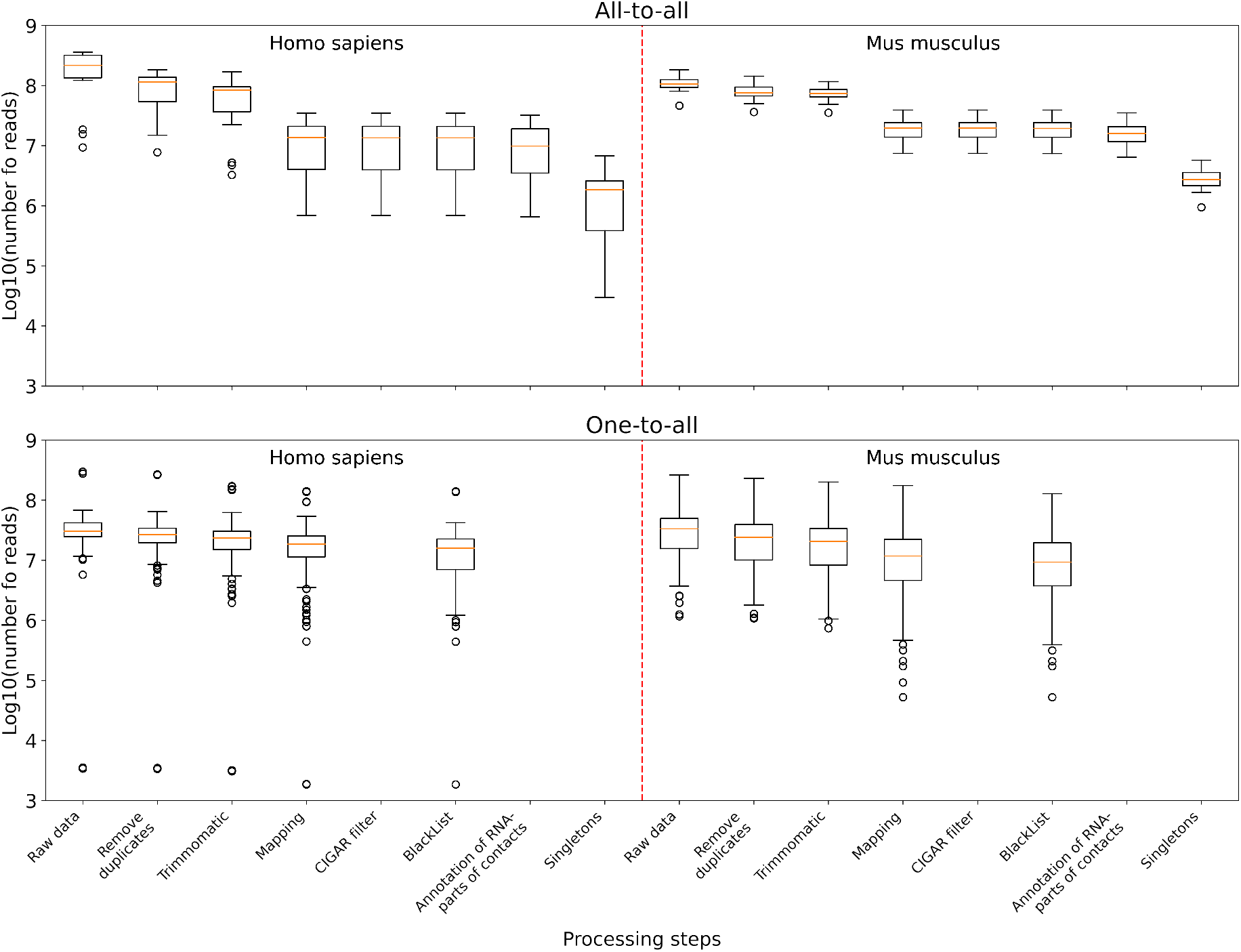
The distribution of the number of reads in the data sets left after the corresponding processing step and all the previous ones. Upper panel: boxplots plotted from all-to-all data, namely 17 human data sets and 18 mouse data sets. Lower panel: boxplots plotted from one-to-all data, namely 159 human data sets and 291 mouse data sets.

## Results

### Database statistics

In humans, 17 all-to-all data sets and 159 one-to-all data sets for 24 RNAs were collected (9 and 53 experiments in the database, respectively). In mice, 18 all-to-all data sets and 291 one-to-all data sets for 31 RNAs were collected (7 and 118 experiments in the database, respectively) (Supplementary Table 2). Since negative controls were not available for all experiments, they were not included in the universal data processing protocol and therefore in the database. In summary, the RNA-Chrom database contains more than 5 billion RNA–chromatin contacts and 232’870 human and 88’914 mouse genes.

### The RNA-Chrom database functionality

With RNA-Chrom, a user can perform two types of analysis of the RNA–chromatin interactions data in real time that can transition into each other. We called the first type of analysis “from RNA”, since the first step is to select the RNA of interest. This analysis allows a user to answer the question “Where does the selected RNA contact chromatin?”. While the second type of analysis, “from DNA”, begins with the selection of the genomic locus of interest, and a user will receive an answer to the question “What RNAs contact with the selected target locus?”. These types of analysis include (Figure 4):

**Fig. 4.**
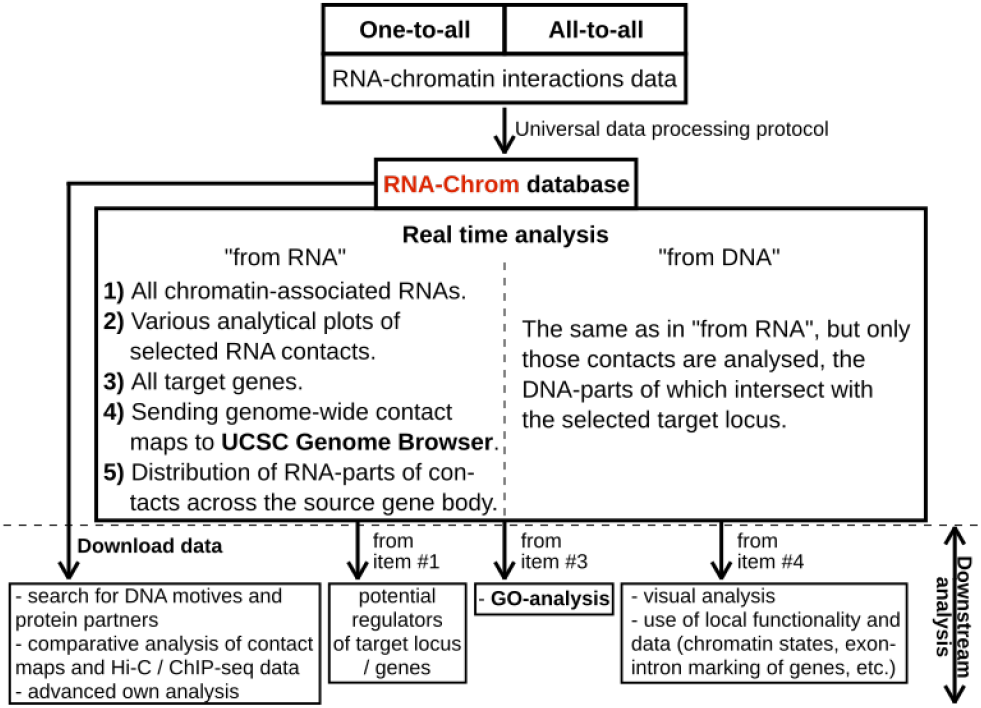
RNA-Chrom functionality and downstream analysis.

1. tables of RNAs contacting the entire genome (“from RNA” analysis) or the selected gene or locus (“from DNA” analysis), with the corresponding characteristics of their contactability;
2. tables of genes with which the selected RNA contacts directly or in the vicinity of 50’000 nucleotides;
3. three types of analytical plots:
  a. contacts density distribution on target locus or on the whole genome;
  b. change in contact density depending on the distance between the RNA source gene and chromatin target loci (“scaling”);
  c. distribution of RNA-parts of contacts across their source gene body.
4. the ability to view contact maps in the UCSC Genome Browser.

In addition, the RNA-Chrom database allows a user to download all pre-processed RNA–chromatin interactions data for a user’s own research or downstream analysis.

### Use case

#### “From RNA” analysis

Our gene annotation consists of 232’870 human and 88’914 mouse genes. Using the web interface, a user can analyse the contacts of any RNA. To perform “from RNA” analysis, the following steps should be taken (Figure 5):

**Fig. 5.**
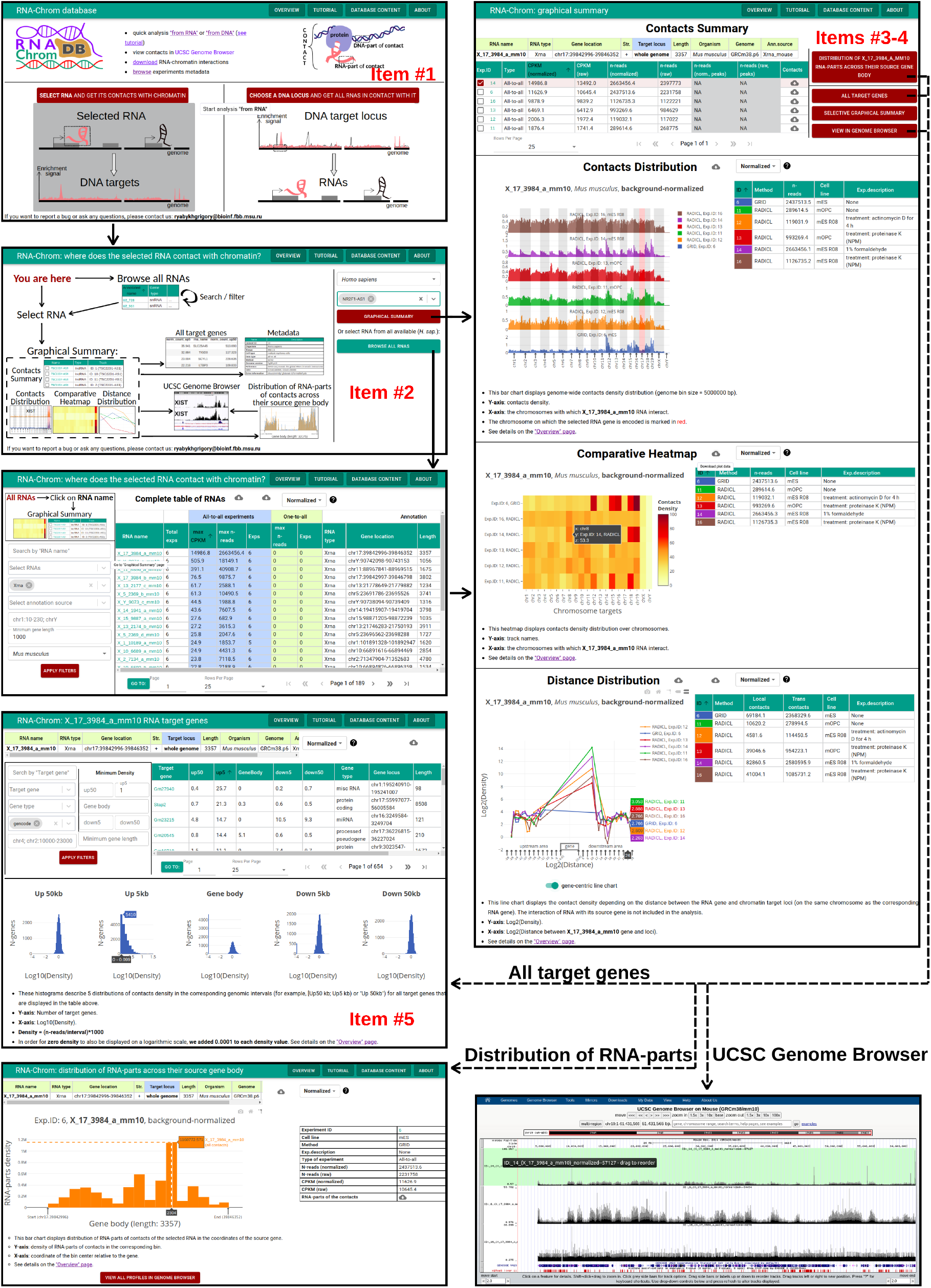
“From RNA” analysis for X_17_3984_a_mm10 RNA (*Mus musculus*). Items #1-5 correspond to the items from the “From RNA analysis” section.

1. A user should choose “from RNA” analysis on the start page. The page “RNA-Chrom: where does the selected RNA contact with chromatin?” will open in a new tab.
2. The next step depends on whether a user knows the RNA name they are interested in or wants to look at the “Complete table of RNAs” (with contact metrics) and select one from it for further analysis.
  a. For example, if a user is interested in NR2F1-AS1 RNA (*Homo sapiens*), they should enter the RNA name in the “Select RNA” field, select the needed RNA in the drop-down list and then click on the “GRAPHICAL SUMMARY” button. The “Graphical Summary” page will open in a new tab that will contain analytical interactive plots and various additional options for further analysis.
  b. A user may click on the “BROWSE ALL RNAS” button. At the “Complete table of RNAs” different filters can be used, such as “Search by RNA name”, “Select RNA names”, “Select RNA types”, “Genomic loci” etc. As an example, a user can fill the filters “Select RNA types”, “Minimum gene length” and “Organism” with the values “Xrna”, “1000” and “*Mus musculus*”, respectively, and then click on the “APPLY FILTERS” button. To go to the “Graphical Summary” page, a user should click on the RNA name they are interested in (for example, X_17_3984_a_mm10, since this RNA has the largest “CPKM”).
3. The “Graphical Summary” page consists of the “Contacts Summary” and three analytical plots: “Contacts Distribution”, “Comparative Heatmap” and “Distance Distribution” (the details can be seen on the “Overview” page).
4. In the “Contacts Summary” table a user can choose, as an example, “Exp.ID: 14” (RADICL, mES R08) and then click on one of the four buttons, for example, “ALL TARGET GENES”. The “All target genes” page will open in a new tab.
5. From the “All target genes” page a user can continue the analysis in “from DNA” mode that will be described below and see what RNAs interact with this target gene, or use filters to get a list of genes that can be downloaded for downstream analysis, such as Gene Ontology.

#### “From DNA” analysis

This type of analysis allows a user to find all RNAs that contact with the selected gene or locus. To perform “from DNA” analysis, the following steps should be taken (Figure 6):

**Fig. 6.**
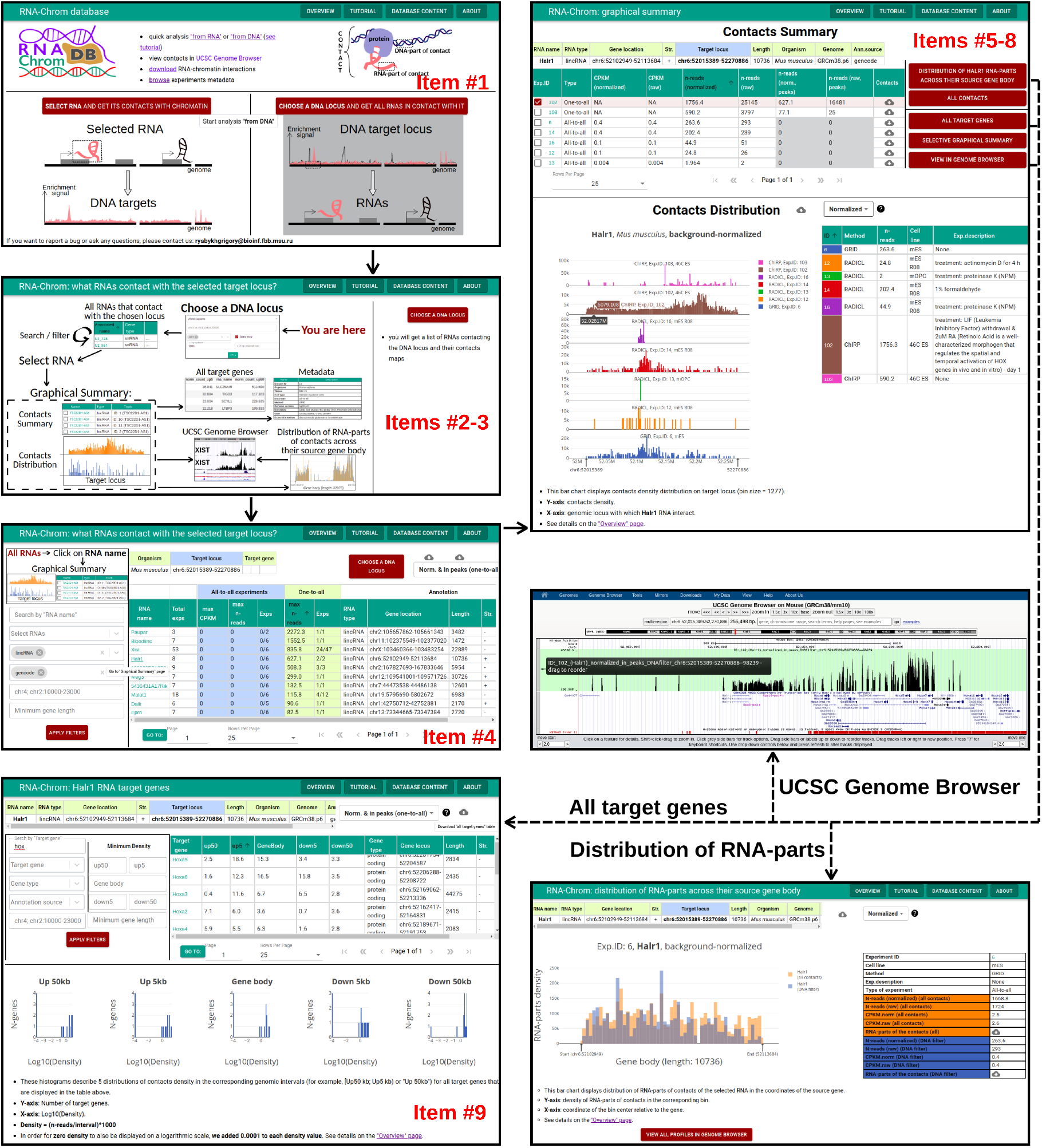
“From DNA” analysis for HoxA-cluster (chr6:52015389-52270886, *Mus musculus*). Items #1–9 correspond to the items from the “From DNA analysis” section.

1. A user should choose “from DNA” analysis on the start page. The page “RNA-Chrom: what RNAs contact with the selected target locus?” will open in a new tab.
2. A user should click on the “CHOOSE A DNA LOCUS” button located on the right side of the page.
3. They can select the organism “*Mus musculus*”, enter the approximate coordinates of the HoxA-cluster (chr6:52015389-52270886) and click on the “APPLY” button. Another way is to select locus by gene name.
4. After locus selection, a list of RNAs that contact with the selected locus appears. To work with this list, different filters can be used, and the list can be sorted in different ways.
  a. As an example, a user can fill the filters “Select RNA types” and “Select annotation source” with the values “lincRNA” and “gencode”, respectively, and click on the “APPLY FILTERS” button.
  b. They can choose normalization “Norm. & in peaks (one-to-all)” and sort the table by the column “max n-reads” (one-to-all).
  c. To go to the “Graphical Summary” page, one should click on the RNA name they are interested in (for example, Halr1, since this RNA is at the top of the table and is also known to be involved in modulating HoxA induction [57]).
5. The “Graphical Summary” page consists of the “Contacts Summary” and the “Contacts Distribution” analytical graph (the details can be seen on the “Overview” page).
6. In the “Contacts Summary” table one can choose, for example, “Exp.ID: 102” (ChIRP, 46C ES, treatment: LIF withdrawal & 2uM RA - day 1).
7. A user may choose normalization “Norm. & in peaks (one-to-all)” and click on the “VIEW IN GENOME BROWSER” button. UCSC Genome Browser will open in a new tab.
8. If one clicks on the “ALL CONTACTS” button, the “Graphical Summary” page corresponding to the “from RNA” analysis will open in a new tab. Here a user can continue the analysis in “from RNA” mode.
9. A user can click on the “ALL TARGET GENES” button, choose normalization “Norm. & in peaks (one-to-all)” and fill the filter “Search by target gene” with the value “hox”. Then they should click on the “APPLY FILTERS” button. As expected, a user will see a lot of Hoxa-genes. From the “All target genes” page one can continue the analysis in “from DNA” mode and see what RNAs interact with this target gene, or download the list of genes for downstream analysis.

#### Content, metadata, overview, tutorial

The “Content” page is a table with complete meta-information for all experiments from the RNA-chrom database (Supplementary Table 2). Here a user can download data for each experiment (contacts with all normalizations, singletons, peaks, etc.). To find out detailed information o n a particular experiment, a user should click on the corresponding “Exp.ID” and go to the “Metadata” page.

The “Metadata” page contains all the metadata for a particular experiment, summary statistics on data processing, “Shares of different RNA types in the total number of contacts” and “Distribution of the number of RNAs according to the number of contacts with the genome” plots. A user can open the “Metadata” page by clicking on the experiment ID everywhere the experiment ID appears, such as on the “Graphical Summary”, “All target genes”, and other pages.

The “Overview” page describes the contents of the RNA-Chrom database and its functionality in detail, specifically what functions are available to a user, what formulas were used to preprocess data for plots and tables, and what information we can extract from those plots and tables.

The “Tutorial” page contains “Basic” and “Advanced” examples of both “from RNA” and “from DNA” assays.

In order to go to the “Content”, “Overview” or “Tutorial” pages, a user should click on the corresponding buttons in the header of the website.

## Discussion

RNA-Chrom is the first manually-curated database that contains a comprehensive collection of genome-wide human and mouse RNA–chromatin interactions data: 16 all-to-all experiments and 171 one-to-all experiments, totaling more than 5 billion RNA–chromatin contacts. We paid special attention to the outstanding procedure of data processing and a user has the opportunity to evaluate the data quality. RNA-Chrom also provides a user-friendly web interface and two types of data analysis (“from RNA” and “from DNA”) which can be used for research.

In order to determine the functional role of RNA in the corresponding DNA locus, additional genome-wide data and annotations are needed, for example, on the structure of chromatin, gene expression, or the localisation of DNA-binding and chromatin-modifying proteins. RNA-Chrom provides a variety of information about the interaction of RNA with chromatin, which can be used in a comparative analysis with other data or as a target for experimental refinements (Figure 4).

In the future, we plan to develop the database by adding new experiments and expanding the list of organisms, as well as to realise several additional normalization procedures that will take into account dependance of contact density on distance between the RNA source gene and chromatin target loci (“scaling”), RNA expression level etc. As we can see, a significant fraction of reads was filtered out due multiple mapping. We are going to develop approaches to the multiple mapping problem in this data. It is expected that the RNA-Chrom database will allow researchers to reach a more systematic level of work with the RNA–chromatin interactome, which will help to expand the understanding of the biological role of non-coding RNAs in a variety of processes.

## Acknowledgments

The authors thank the anonymous reviewers for their valuable suggestions. This work is supported by RFBR grant #20-04-00459 A.

## Footnotes

1 https://nodejs.org/en/

2 https://reactjs.org/

3 https://github.com/reduxjs/redux

4 https://mui.com

5 https://plot.ly/javascript/

6 https://pgjones.gitlab.io/quart/

7 https://clickhouse.com/

8 http://www.ncbi.nlm.nih.gov/geo/

9 https://www.ebi.ac.uk/ena/browser/home

