## Supplementary Text 1, Supplementary Text 2, Supplementary Figure 1 for "RNA-Chrom: a manually-curated analytical database of RNA–chromatin interactome"

#### Remove duplicates

Possible PCR duplicates of RNA-DNA pairs of reads (all-to-all data) and paired-end reads (one-to-all data) were removed via FastUniq tool [1], while SeqKit rmdup [2] was used to process single-end reads (one-to-all data). For iMARGI [3, 4] data sets we followed the original paper authors' recommendations to perform this step after "Add restriction site".

It is important to note that all replicas were processed independently starting from the "Remove duplicates" step and up to "BlackList".

#### Add restriction site

Since endonucleases were used to fragment genomic DNA in the GRID-seq [5], Red-C [6] and iMARGI [3, 4] methods (sonication was used in RADICL-seq [7]), it was important to filter out reads in which one of the ends did not end with half of the restriction site. Further, the second part of the restriction site was added to the corresponding ends of the reads, which slightly increased the length of the reads and increased the efficiency of unique mapping. This procedure was performed in strict accordance with the recommendations of the original articles.

#### Trimmomatic

We used TRIMMOMATIC (v0.39) [8] ( parameters: `window size = 5, quality threshold = 26, minlen = 14` for detection of low-quality position in each forward and reverse read. Few low-quality cases with more than 50% discarded data after trimming (Supplementary Table 3) were re-trimmed with a more permissive "quality threshold" of 22 and an additional `LEADING:22` parameter.

#### Mapping

One-to-all data were mapped to the genome with HISAT2 program (version 2.1.0) [9] (parameters for single-end reads: -k 100 –no-spliced-alignment –no-softclip; for paired-end reads: `-k 100 --no-spliced-alignment --no-softclip --no-discordant --no-mixed`). DNA-parts and RNA-parts of the contacts (all-to-all data) were independently mapped by the same program (parameters for DNA-parts: `-k 100 --no-spliced-alignment --no-softclip`, for RNA-parts: `-k 100 --no-softclip --dta-cufflinks --known-splicesite-infile`). Splice site annotations for the respective genomes were obtained using the "hisat2_extract_splice_sites.py" script [9]. SAM files were filtered for unique mappings with at most 2 mismatches relative to the reference genome.

#### CIGAR

Reads corresponding to RNA-parts can be mapped in three ways:

1. with a complete match with the reference genome along the entire length of the read (CIGAR of the "25M" type, where M – match). Such reads went on without changes;
2. containing one missing interval (CIGAR of the form "30M65N10M", where M – match, N – skipped region). For such reads, the longest section mapped without breaks was left;
3. more complex mapping options (reads with complex splicing): multiple missing intervals (CIGAR of the form: "8M1113N56M79N8M"), mapping with insertions or deletions. All such reads were removed.

#### BlackList

DNA-parts of the contacts that fell into the regions from the ENCODE BlackList for GRCh38 and GRCm38 (accession: ENCSR636HFF) were removed. For one-to-all data, "input" (data with background nonspecific contacts) was not filtered by the BlackList to avoid edge effects.

#### Annotation of RNA-parts of contacts

If the gene names were repeated in the general gene annotation (more details about gene annotations can be found in the relevant section here: `https://rnachrom2.bioinf.fbb.msu.ru/protocol`, then a serial number was assigned to them so that all gene names in the database are unique. For example, the "WASIR1" gene was found twice in the gencode annotation, so we assigned "WASIR1_1" and "WASIR1_2" to the copies, respectively. Since one-to-all data does not have any information on the RNA-parts of the contacts, we assigned to each DNA-part from the experiment the coordinates of the corresponding RNA source gene studied in the corresponding experiment. In the case of all-to-all data, if the RNA-part of the contact intersects a gene by at least 1 nucleotide, this RNA-part was assigned to this gene. If the RNA-part of the contact intersects more than one gene at the same strand, this RNA-part was assigned to the gene showing the highest coverage by RNA-parts, which was determined as the total number of RNA-parts mapped to the gene normalized to the gene length.

Note that in this and subsequent steps, the replicas are already merged to increase the amount of data and coverage.

#### X-RNAs assembly

A substantial amount of RNA-parts was not annotated by any of the used gene annotations. Some of these unknown parts may belong to unknown ncRNAs. Transcripts not corresponding to any known gene (from GENCODE database [10]: annotation version 35 for human and 25 for mice) were assembled using StringTie [11] and then filtered by several criteria, e.g. length, distance to the closest known gene on the same strand, conservation on different taxonomic levels and high coverage. We called the group of transcripts that passed all the filters "X-RNAs". To the X-RNAs the unique IDs were assigned based on their genome location. E.g. X_1_13_a_hg38 is the X-RNA located on chromosome 1 of the human genome (version hg38) in the 13th bin (each chromosome was divided into bins of 10'000 bp). And the letter "a" indicates that the source gene of this X-RNA is the first in the bin relative to the beginning of the corresponding chromosome.

#### Contacts normalization

For each experiment, the following procedure was carried out. A single value ("n-reads (raw)" = 1) was assigned to each contact, since a single contact corresponds to 1 pair of RNA- and DNA-parts. After that, we defined background contacts depending on the type of data:

1. **All-to-all experiments.** According to the approach proposed in the GRID-seq [5] protocol, we defined background contacts as the total number of mRNAs trans-contacts with each genomic site (not with the parental chromosome). The genome was divided into 500 bp bins, and for each bin we summed up the number of trans-contacts made with this bin by protein-coding mRNAs (the 50 most contacting and 1000 least contacting mRNAs were removed for each experiment).

2. **One-to-all experiments.** The genome was divided into 500 bp bins, and for each bin we summed up the number of "input" contacts whose centers fell into the corresponding bin (if there was no "input" library for a particular experiment, then the background was made constant, that is, we assigned exactly one contact to each bin).

Then we smoothed the obtained signal with StereoGene (v.2.20) [12] (parameters: `bin = 500`, `wSize = 1000000`, `flankSize = 10000`, `kernelSigma = 3000`, `kernelType = NORMAL`) and used it as a background signal – "n-reads (background)" or $n_{bg}$.

Then we normalized each $n_{raw}$ by the value of the background signal ($n_{bg}$) in the genomic coordinate where the DNA-parts were mapped. To work with DNA-parts mapped to the regions with zero value of the background signal, we added the pseudocount to the $n_{bg}$. And thus, we obtained nomalized value ("n-reads (normalized)" or $n_{norm}$). This normalization ensures that the sum of the normalized values is equal to the number of reads in the experiment (Eq.1).

$$n_{norm} = \frac{n_{raw}}{n_{bg} + 0.5} \cdot (\sum n_{raw} \ / \ \sum \frac{n_{raw}}{n_{bg} + 0.5}) \qquad (1)$$

For one-to-all experiments, the peaks (a genome regions enriched in contacts of the RNA with chromatin) were additionally called using the MACS2 [13] program with the following parameters: `-Q 0.05 -FORMAT BED` (if single-end) or `-Q 0.05 -FORMAT BEDPE` (if paired-end). Contacts, the DNA-parts of which intersected the peaks by at least 1 bp, will be used in the further construction of analytical plots with previously defined "n-reads (raw)" and "n-reads (normalized)". Applying this filter, we obtained "n-reads (raw & in peaks)" and "n-reads (norm. & in peaks)".

#### Annotation of DNA-parts of contacts

For each gene we calculated the following five intervals: gene body, 5 Kb upstream and 5 Kb downstream from the gene, from 5 to 50 Kb upstream and from 5 to 50 Kb downstream from the gene according to the gene strand. Hence, the intervals don't intersect. DNA-parts of the contacts were annotated by all the intervals. The coordinates of the DNA-part and the corresponding intervals must intersect by at least 1 nucleotide.

### Supplementary Text 2

During the data processing steps we noticed that RNA-parts of the contacts in a number of experiments could represent not those parts of genes sequences, from which the corresponding RNAs were transcribed, but reverse complements of those sequences. In other words, In some experiments, the "forward" strand of the cDNA read part could be sequenced, and in others — the "reverse" one.

In order to determine whether that hypothesis was true, an experiment was carried out based on the following assumption: in any viable cell line, the genes of ribosomal proteins must be highly expressed, and it is likely that a significant part of the data we have is precisely the messenger RNAs of these proteins contacting with chromatin on their way to nuclear pores. For each data set, we can select the RNA-parts of the contacts that were aligned within the coordinates of the genes of ribosomal proteins on both chains, and then calculate the fractions of reads aligned on the gene chain and the chain complementary to the gene (Figure 2).

If more reads were mapped to the gene chain than to its complementary one, then during sequencing, the "correct" cDNA chain was read corresponding to the sequence of RNA in contact with chromatin, and vice versa. The RNA-parts from experiments that have "wrong" cDNA chain sequences needed to be reversed before future analysis, although it was not obviously stated in any of the original papers.

Notably, when looking at human data, the following strands were read: the "correct" cDNA strand (in the case of the Red-C [6] experiment); the reverse strand (for GRID-seq [5] and iMARGI [3, 4] experiments). Whereas in the case of MARGI [14], it seems that mostly a random strand was read, and the orientations of the RNA-parts of the contacts were lost (Figure 2). It can be seen that for some MARGI data sets (SRR5278097, SRR5278097, SRR5278100, SRR5278102) strands were uniquely determined. However, due to the low gene coverage of ribosomal proteins in these data sets (Supplementary table 4) and to the loss of RNA-parts orientations in other MARGI data sets, we decided to exclude the MARGI experiment from any further analysis.

Looking at mouse data, we can see that the mouse GRID-seq behaves like a human GRID-seq, and RADICL-seq [7] behaves like Red-C (Figure 3).

### Supplementary Tables (see file "RNA-Chrom supplementary tables")

1. **Table 1.** Data processing statistics.
2. **Table 2.** Database content.
3. **Table 3.** 28 data sets that have been re-trimmed with a more permissive "quality threshold" of 22 and an additional LEADING:22 parameter.
4. **Table 4.** Data that are involved in clarifying the orientation of the RNA-parts of human contacts.

### Supplementary Figures

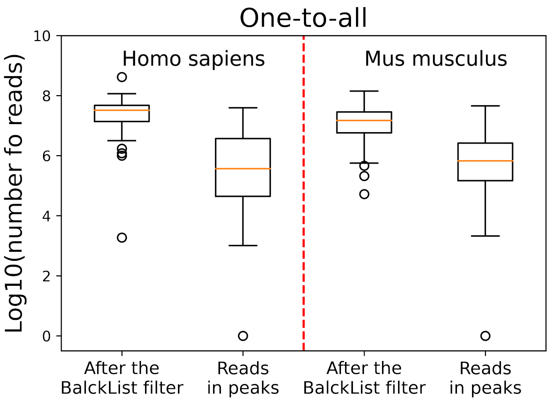


**Fig. 1.** Distribution of the number of reads (contacts) in one-to-all experiments (the replicas are already merged), remaining after all processing steps ("BlackList" is the last step) and after additional filtering: the intersection with MACS2 peaks ("Reads in peaks"). In order for data sets with zero "Reads in peaks" to also be displayed, we added 1 read to each experiment.

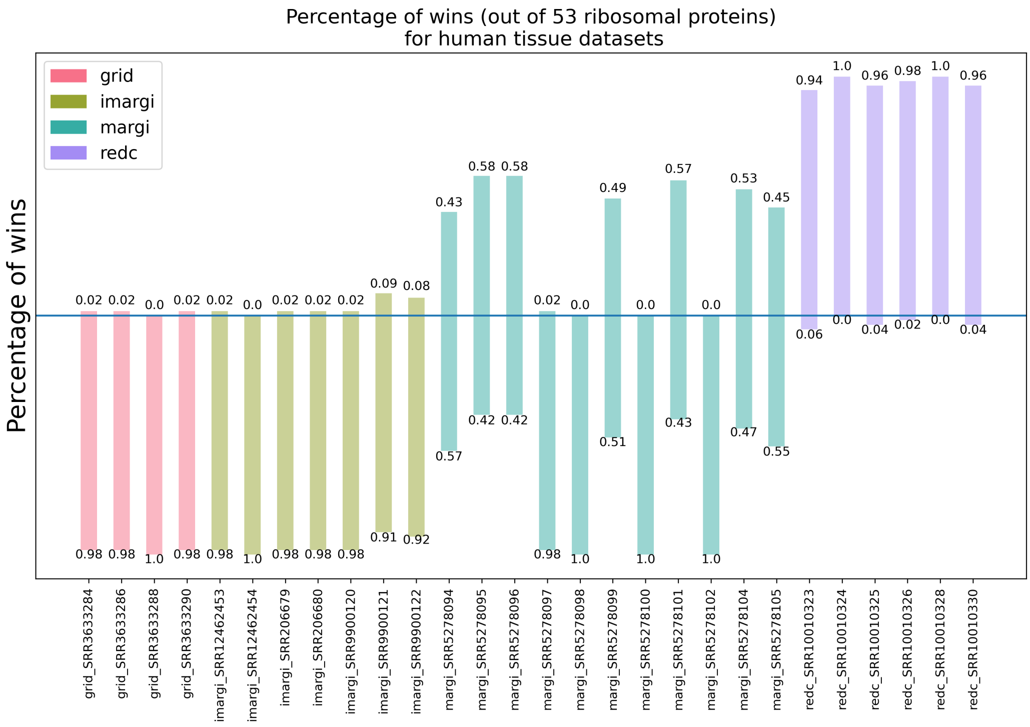


**Fig. 2.** Estimation of orientation of RNA-parts of the contacts for human data sets on a subsample of 53 ribosomal protein-coding genes (RPL22, RPL11, RPL5, RPL31, RPL37A, RPL32, RPL15, RPL14, RPL29, RPL24, RPL22L1, RPL39L, RPL35A, RPL9, RPL34, RPL37, RPL26L1, RPL10A, RPL7L1, RPL7, RPL30, RPL8, RPL35, RPL12, RPL7A, RPLP2, RPL27A, RPL41, RPL6, RPLP0, RPL21, RPL10L, RPL36AL, RPL4, RPLP1, RPL3L, RPL13, RPL26, RPL23A, RPL23, RPL19, RPL27, RPL38, RPL28, RPL3, RPL36A, RPL13A, RPL39, RPL10 , RPL17, RPL36, RPL18A, RPL18).

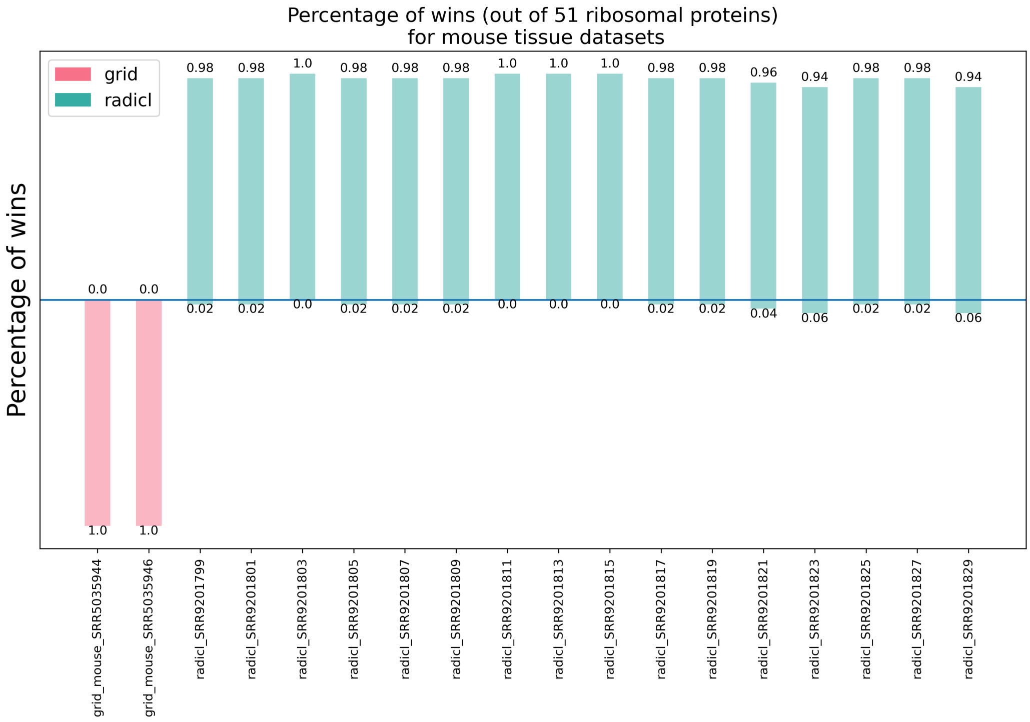


**Fig. 3.** Estimation of orientation of RNA-parts of the contacts for mouse data sets on a subsample of 51 ribosomal protein-coding genes (Rpl7, Rpl31, Rpl37a, Rpl7a, Rpl12, Rpl35, Rpl22l1, Rpl34, Rpl11, Rpl22, Rpl9, Rpl5, Rplp0, Rpl6, Rpl21, Rpl32, Rpl28, Rpl13a, Rpl18, Rpl27a, Rplp2, Rpl18a, Rpl13, Rplp1, Rpl4, Rpl29, Rpl14, Rpl41, Rpl26, Rpl23a, Rpl23, Rpl19, Rpl27, Rpl38, Rpl10l, Rpl15, Rpl37, Rpl30, Rpl8, Rpl3, Rpl39l, Rpl35a, Rpl24, Rpl3l, Rpl10a, Rpl7l1, Rpl36, Rpl17, Rpl39, Rpl10, Rpl36a).
